# Cardiolipin Inhibits the Noncanonical Inflammasome by Preventing LPS Binding to Caspase 4/11 to Mitigate Endotoxemia in Vivo

**DOI:** 10.1101/2024.10.14.618051

**Authors:** Malvina Pizzuto, Mercedes Monteleone, Sabrina Sofia Burgener, Jakub Began, Melan Kurera, Jing Rong Chia, Emmanuelle Frampton, Joanna Crawford, Monalisa Oliveira, Kirsten M. Kenney, Jared R. Coombs, Masahiro Yamamoto, Si Ming Man, Petr Broz, Pablo Pelegrin, Kate Schroder

## Abstract

In Gram-negative bacterial sepsis, excessive caspase 4/11 activation in response to circulating bacterial lipid LPS (endotoxemia) can cause organ damage and mortality. Current inhibitors of caspase 4/11 also block caspase 1 activity and are therefore not appealing clinical candidates for treating Gram-negative sepsis. Here, we identify double-unsaturated 18:2 cardiolipin as a selective inhibitor of caspase 4/11-dependent inflammatory cytokine secretion and pyroptosis, without affecting caspase-1 responses. Cardiolipin targets the CARD domain of caspase 4/11, impeding its interaction with LPS to restrain caspase 4/11 activation, thereby suppressing endotoxemia-induced systemic inflammation *in vivo*. Thus, we present cardiolipin as a promising candidate for preventing endotoxemia-induced sequelae in sepsis while preserving caspase-1-driven anti-microbial immune responses. By identifying cardiolipin as a specific caspase 4/11 inhibitor, we provide an urgently-needed tool for studying caspase 4/11 functions in inflammatory pathways, and open the way to studies of noncanonical inflammasome regulation by endogenous cardiolipin.

## Introduction

Sepsis is a life-threatening condition involving organ dysfunction caused by a dysregulated host response to infection (Singer et al., 2016). Sepsis can lead to septic shock, where vital organs are damaged, accounting for 19% of global deaths (Cecconi et al., 2018; Fleischmann et al., 2016; Giamarellos-Bourboulis et al., 2024; Rudd et al., 2020).

Over 100 Phase II and Phase III clinical trials have unsuccessfully attempted to treat sepsis by generic immunosuppression or by targeting endogenous inflammatory mediators such as tumour necrosis factor (TNF) or interleukin (IL)- 1β (Giamarellos-Bourboulis et al., 2024; Marshall, 2014). Recent mechanistic studies have elucidated the pathways that cause sepsis-induced organ damage, opening new avenues for more specific and effective treatments (Cheng et al., 2017; Deng et al., 2018; Wei et al., 2024). 30-50% of total sepsis cases are caused by Gram-negative bacteria (Alberti et al., 2002; Martin et al., 2003); here, activation of the inflammatory cascades by high concentrations of circulating bacterial lipopolysaccharide (LPS) cause endotoxemia, organ damage and lethality (Gabarin et al., 2021; Opal and Glück, 2003). Endotoxemia is also observed in viral infections that damage the intestinal barrier (e.g., SARS-CoV- 2), causing LPS to enter the circulation (Assimakopoulos et al., 2022). Unrestrained LPS signalling causes cell death and systemic inflammation characterised by the induction of proinflammatory cytokines, such as TNF and IL- 1β (Gabarin et al., 2021), which contribute to tissue injury (Takahama et al., 2024).

Extracellular LPS binding to Toll-like Receptor 4 (TLR4) activates various transcription factors, including NF-κB, leading to the production and release of inflammatory cytokines, including TNF and IL-6 (Gay et al., 2014; Kawai and Akira, 2010; Park et al., 2009). TLR4 activation also induces the expression of proteins involved in the inflammasome pathways, including nucleotide-binding domain leucine-rich repeat and pyrin domain-containing protein 3 (NLRP3), absent in melanoma 2 (AIM2), caspase 11 (CASP11), and pro-IL-1β (Schroder and Tschopp, 2010). This poises cells to respond to microbial threats by activating inflammasomes to produce cytokines (e.g., IL-1β, IL-18) essential for antimicrobial defences (Schroder and Tschopp, 2010).

Canonical inflammasomes are large cytosolic signalling complexes composed of a sensor protein (e.g. NLRP3 or AIM2) that detects microbial or danger signals, the adaptor protein apoptosis-associated speck-like protein containing a caspase recruitment domain (ASC), and the effector protein caspase 1 (CASP1) (Schroder and Tschopp, 2010). NLRP3 is activated by a wide range of stress signals, including potassium efflux inducers such as nigericin (Broz and Dixit, 2016). NLRP3 activation induces the assembly of the canonical inflammasome complex and the activation of CASP1. Active CASP1 cleaves pro-IL-1β, pro-IL-18 and gasdermin D (GSDMD) into their active, pro-inflammatory forms — IL-1β, IL-18, and N-terminal GSDMD (Nt-GSDMD) (Chan and Schroder, 2020). Subsequently, Nt-GSDMD forms pores in the cell membrane to release IL-1β and IL-18, and induces a lytic form of cell death known as pyroptosis (Orning et al., 2019). Pyroptosis ejects intracellular contents such as lactate dehydrogenase (LDH) and pro-inflammatory damage-associated molecular patterns (DAMPs) into the extracellular space (Broz et al., 2019).

The NLRP3 inflammasome can also signal downstream of the activation of the intracellular LPS sensors – caspase 4 in humans and caspase 11 in mice (hereafter CASP4/11) – through the so-called noncanonical inflammasome pathway (Kayagaki et al., 2011; Schmid-Burgk et al., 2015). Upon binding to LPS, CASP4/11 dimerises and auto-cleaves to form an active CASP4/11 species, which cleaves GSDMD to induce pyroptosis (Chan et al., 2023; Kajiwara et al., 2014; Lee et al., 2018; Ross et al., 2018). Here, GSDMD pores trigger potassium efflux that activates the NLRP3/CASP1 inflammasome, causing IL-1β and IL-18 processing and release and the amplification of the inflammatory response (Baker et al., 2015; Kayagaki et al., 2015; Schmid-Burgk et al., 2015; Viganò et al., 2015).

Clinical trials that reduced LPS-induced inflammation (e.g. the TLR4 antagonist eritoran, broad-acting immunosuppressants, blockers of TNF or CASP1/IL-1β signalling) were unsuccessful (Dinarello, 2018; Marshall, 2014), perhaps because these also blocked anti-microbial defence. Indeed, immunosuppressed patients who cannot mount a canonical NLRP3/CASP1 inflammasome response have the highest sepsis mortality rates (Martínez-García et al., 2019). This suggests that indiscriminate targeting of LPS signalling in a manner that also blocks other immune surveillance pathways is detrimental to patients, highlighting the urgent need for more specific immunomodulators.

Like TLR4 and CASP1, CASP4/11 signalling contributes to host protection against infection. LPS-induced CASP4/11 activation in macrophages, epithelial and other cells is vital to detect bacterial threats and mount an antimicrobial immune response (Aachoui et al., 2015, 2013; Chen et al., 2018; Enosi Tuipulotu et al., 2023; Knodler et al., 2014; Kobayashi et al., 2013; Kovacs et al., 2020; Kumari et al., 2021; Wang et al., 2018). However, the extremely high LPS concentrations reached during Gram-negative bacterial sepsis and endotoxemia can cause aberrant CASP4/11 activation (Chen et al., 2019; Cheng et al., 2017; Deng et al., 2018; Hagar et al., 2013; Kajiwara et al., 2014; Kayagaki et al., 2015, 2011; Tang et al., 2018; Wei et al., 2024). In preclinical models of sepsis and endotoxemia, including a transgenic model of human CASP4 expression in mice, signal blockade of CASP11 or CASP4, protected mice from organ damage and death (Cheng et al., 2017; Deng et al., 2018; Hagar et al., 2013; Kayagaki et al., 2013; Wei et al., 2024). While such reports highlight the potential benefit of specific inhibition of CASP4/11 for treating sepsis, such CASP4/11-specific inhibitors remain to be identified.

Cardiolipin is a tetra-acylated lipid present in mitochondrial and bacterial membranes. Research by us and others showed that double-unsaturated cardiolipin with 18 carbon atom chain length (hereafter called CL) inhibits TLR4 (Balasubramanian et al., 2015; Pizzuto et al., 2019; Wenzel et al., 2021). CL occupies the LPS binding site of TLR4 to prevent LPS ligation and specifically inhibits TLR4 without affecting signalling by other TLRs (Pizzuto et al., 2019). Similar to CL, the TLR4 antagonists eritoran and its analogue LPS derived from *R. sphaeroides* (RS-LPS) are also lipids that block the LPS binding site on TLR4, but possess only a single mono-unsaturated chain. The capacity of CL, eritoran and RS-LPS to inhibit other LPS receptors has not been tested.

Here we examined the capacity of CL and RS-LPS to inhibit CASP4/11 activation. We found that CL, but not RS-LPS, is a CASP4/11 inhibitor. Mechanistically, CL targets the CASP4/11 CARD domain and prevents LPS binding and resultant CASP4/11 signalling. CL specifically inhibit CASP4/11 over CASP1 and suppresses cell death and inflammatory cytokine secretion *in vitro* and *in vivo*. These findings identify CL as a novel and specific CASP4/11 inhibitor. This discovery enables new approaches for understanding the pathological functions of CASP4/11 in experimental sepsis and offers new avenues for translation to urgently-needed novel treatments for human sepsis.

## Results and Discussion

### Cardiolipin but not RS-LPS inhibits caspase 4/11 signalling

We first sought to determine whether diunsaturated CL or monounsaturated RS-LPS suppressed CASP4/11 noncanonical inflammasome signalling outputs. We treated Pam3CSK4 (Pam)-primed human monocyte-derived macrophages (HMDM) or bone marrow-derived murine macrophages (BMDM) with the noncanonical inflammasome activator LPS from *E. coli* B4, delivered intracellularly using lipofectamine (LTX) or Cholera toxin B (CTB), in the absence or presence of CL. We quantified IL-1β and LDH release as a measure of noncanonical inflammasome signalling induced by intracellular LPS (iLPS). iLPS induced IL-1β and LDH release in HMDM and BMDM, and this was suppressed by CL, but potentiated by RS-LPS (Fig 1 A-B); the latter result is in line with a report that RS-LPS induced LDH and IL-1β release in macrophage (Lagrange et al., 2018). This indicates that CL, but not RS-LPS, dampens noncanonical inflammasome signalling and suggests that possession of only one mono- unsaturated chain is insufficient for lipids (e.g., RS-LPS and its analogue eritoran) to inhibit CASP4/11.

**Figure 1:**
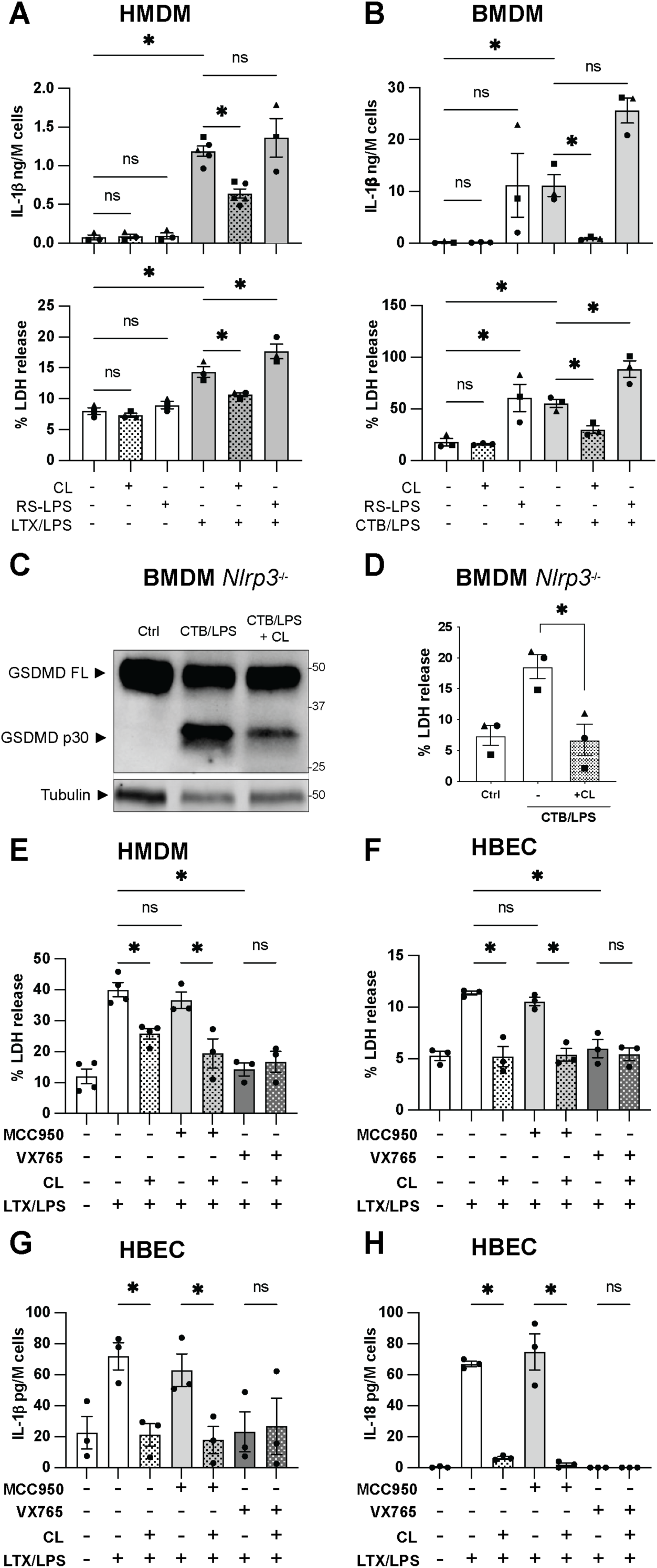
CL but not RS-LPS inhibits noncanonical inflammasome signalling independently of NLRP3 in primed-macrophages and epithelial cells. **A - B** Human Monocytes-Derived Macrophages (HMDM) from healthy donors (A) or Bone Marrow-Derived Macrophages (BMDM) from wild-type (WT) mice (B) were incubated for 4 hours with 1 µg/mL Pam3CSK4. Cell culture medium was replaced with OptiMEM plus HEPES (Ctrl), 10 µM CL, or 5 µg/mL of RS-LPS (complexed with LTX) in the absence or presence of 2 µg/mL of LPS complexed with LTX (A) or CTB (B). Cells were incubated for 4 (A) or 18 hours (B). Cell supernatants were analysed for cleaved IL-1β (ELISA) and LDH (cytotoxicity assay). **C - D** BMDM from *Nlrp3*^-/-^ mice were incubated for 4 hours with 1 µg/mL Pam3CSK4, before cell culture medium was replaced with OptiMEM plus HEPES (Ctrl), or 2 µg/mL of LPS complexed with CTB in the presence of HEPES (-), or 10 µM CL (CL). Cells were incubated for 18 hours. GSDMD and tubulin expression and cleavage were assessed in mixed supernatants and lysates by western blot (C). LDH release was quantified by cytotoxicity assay (D). **E - H** HMDM from healthy donors (E) or Human Bronchial Epithelial Cells (HBEC) (F-H) were incubated for 4 (E) or 18 (F-H) hours with 1 µg/mL Pam3CSK4. Cell culture medium was replaced with OptiMEM with vehicle (DMSO) or 10 µM MCC950 (NLRP3 inhibitor), or 10 µM VX-765 (CASP1/4 inhibitor). Cells were incubated for 1 hour, and then 2 µg/mL of LPS complexed with LTX was added in the presence of HEPES (-) or 10 µM CL (CL). Cells were incubated for 4 hours. Cell supernatants were analysed for cleaved IL-1β (ELISA) and LDH (cytotoxicity assay). **Data information:** Each symbol is the mean of technical triplicates from an independent biological replicate. Bars are the mean of three or more independent biological replicates (n = 3 to 5) ± SEM. **Statistical analysis:** A-B and E-H One-way ANOVA Šídák’s multiple comparisons test. D unpaired t- test. Significant difference for p<0.05 (*), not significant p≥0.05 (ns).

CL was previously reported to activate NLRP3 inflammasome signalling in a macrophage cell line (Iyer et al., 2013; Murphy and O’Neill, 2024). To ensure that inhibition of iLPS signalling was not due to CL-induced NLRP3-dependent cell death, we treated primed HMDM and BMDM with CL alone. In both primed HMDM and BMDM, CL failed to induce the release of IL-1β or LDH (Fig 1 A-B), suggesting that 18:2 CL does not activate inflammasome signalling in primary human and murine macrophages.

We also tested that CL was not toxic to macrophages, by treating primed BMDM or HMDM with escalating doses of CL (from 1 to 50 µM). Cell toxicity was monitored by measuring LDH release and YoPro uptake, all of which confirmed that CL did not induce macrophage cell death (Fig S1 A-C). To determine whether CL inhibits noncanonical inflammasome signalling to a broad repertoire of LPS species, we treated Pam-primed BMDM with intracellular LPS derived from several bacterial strains (*E. coli* K12, *Salmonella minnesota* R595, *Pseudomonas aeruginosa*) as compared to the *E. coli* B4 LPS previously used. CL suppressed LDH and IL-1β release induced by all LPS species (Fig S1 D-E), indicating that CL may inhibit noncanonical inflammasome signalling to a wide variety of bacterial strains.

### Cardiolipin inhibits the noncanonical inflammasome upstream of NLRP3 activation

Given that CASP4/11 induces GSDMD cleavage and resultant NLRP3 inflammasome signalling, we next investigated whether NLRP3 is a p potential target of CL. We blocked NLRP3 signalling in primed BMDM (through *Nlrp3* knockout) or HMDM (by pre-treating cells with the NLRP3 inhibitor MCC950) before treating cells with iLPS in the absence or presence of CL. CL inhibited iLPS-induced GSDMD cleavage and LDH release in *Nlrp3*^-/-^ murine macrophages (Fig 1 C-D) and MCC950-treated HMDM (Fig 1 E). The CASP1/4 inhibitor VX765 blocked iLPS-induced LDH release (Fig 1 E), confirming that HMDM death was pyroptotic. Collectively these data demonstrate that in human and murine macrophages, CL inhibits noncanonical signalling upstream and independently of NLRP3.

In murine macrophages, NLRP3/CASP1 signalling is required for IL-1β cleavage and release downstream of CASP11 activation. In human cells, however, CASP4 can cleave pro-IL-18 and pro-IL-1β independently of NLRP3; accordingly, in human epithelial cells that do not express NLRP3, CASP4 activity is sufficient for release of mature IL-18 and IL-1β (Chan et al., 2023; Shi et al., 2023). To determine whether CL blocks CASP4-mediated pro-IL-1β independently of NLRP3, we thus tested human bronchial epithelial cells (HBEC). CL suppressed iLPS-induced LDH, IL-1β and IL-18 release from HBEC, and this was unaffected by MCC950 as expected (Fig 1 F-H). Verifying that these signalling outputs are inflammasome-dependent, VX765 blocked iLPS-induced LDH, IL-1β and IL-18 release (Fig 1 F-H). Thus, CL blocks signalling by CASP4 in the absence of NLRP3 signalling. Moreover, these data extend the CASP4 inhibitory activity of CL to non-myeloid cells.

### CL binds to the CARD domain of CASP4/11, preventing LPS binding and consequent CASP4/11 activation

To define the mechanism by which CL suppresses noncanonical inflammasome signalling, we initially examined whether CL blocks CASP4/11 intracellular signalling in BMDM and HMDM. Macrophages were left unprimed (UP), or were primed with Pam3CSK4 and then treated with iLPS in the presence or absence of CL. The cell culture medium was precipitated and resuspended in cell lysates for immunoblot analyses of CASP4/11, GSDMD, IL-1β, and CASP1 cleavage. HMDM control samples (unprimed and primed) express CASP4, CASP1 and GSDMD and their expression is not affected by priming, while pro-IL-1β was strongly induced by priming (Fig 2 A). Notably, CL did not affect the expression of these full-length proteins but suppressed iLPS-induced GSDMD, CASP1, and IL-1β cleavage (Fig 2 A). Although we were unable to detect the cleaved fragment of CASP4, iLPS induced a loss of full-length CASP4 protein that was suggestive of cleavage, and this was blocked by co-incubation with CL (Fig 2 A). Similarly, in BMDM, iLPS induced CASP11, IL-1β, GSDMD, and CASP1 cleavage, and this was reduced by CL co-administration without affecting the expression of full-length GSDMD or CASP1 (Fig 2 B). In all, these data indicate that in human and murine macrophages, CL inhibits iLPS-induced CASP4/11 activation and resultant GSDMD, CASP1 and IL-1β cleavage.

**Figure 2:**
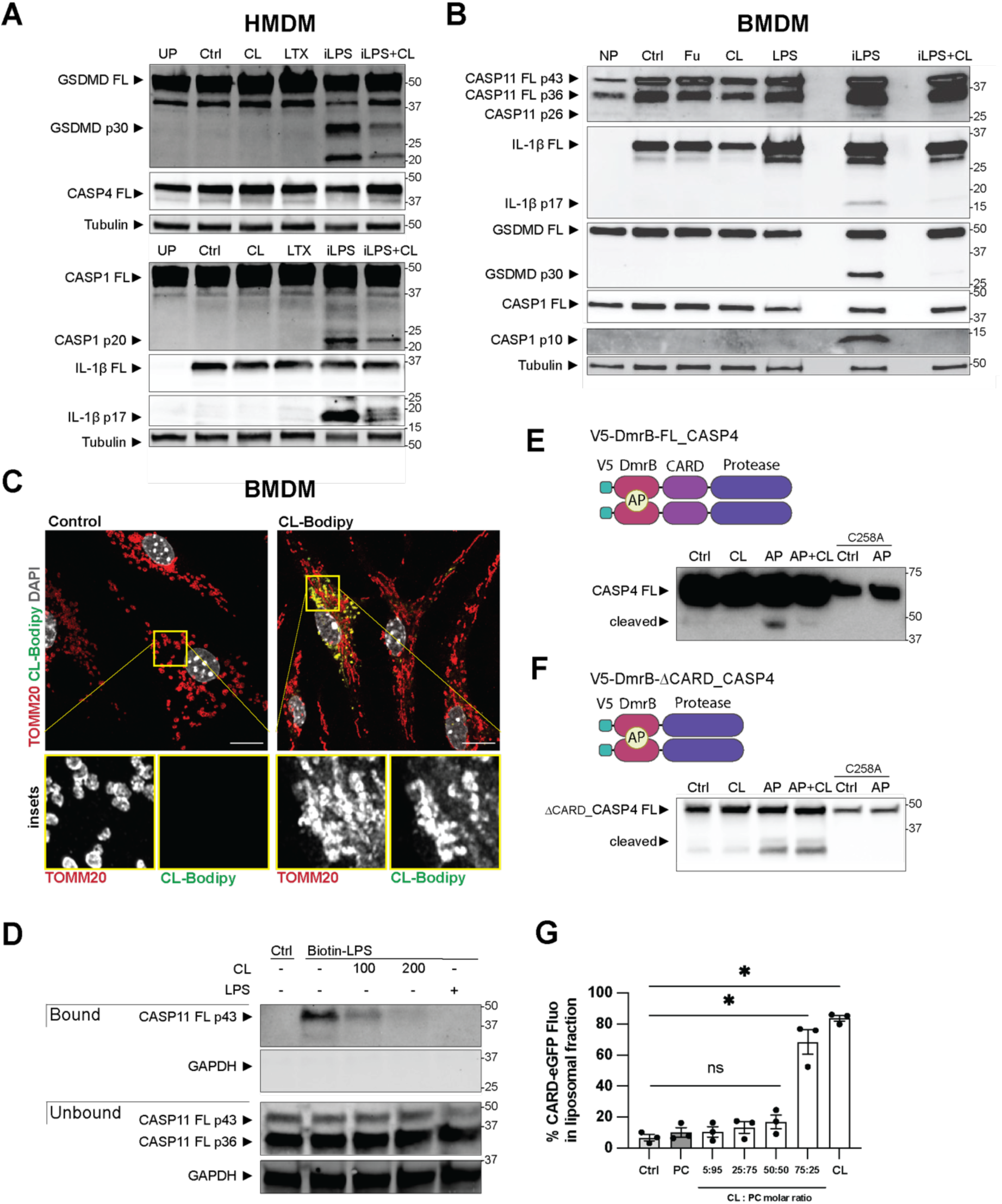
CL reaches the cell interior and binds to the CARD domain of CASP4/11, preventing LPS binding and consequent CASP4/11 activation. **A - B** HMDM (A) or WT BMDM (B) were incubated for 4 hours with cell culture medium (unprimed, NP) or 1 µg/mL Pam3CSK4 (all other conditions). Cell culture medium was then replaced with OptiMEM (Ctrl), FuGENE HD 0.5% v/v, LTX 0.25% v/v, 10 µM CL, or 2 µg/mL of LPS complexed with 0.25% v/v LTX (A) or 0.5% FuGENE HD (B) in the presence of HEPES vehicle (iLPS) or 10 µM CL (iLPS + CL). Cells were incubated for 4 hours. GSDMD, CASP1, CASP4, CASP11, IL-1β, and tubulin expression and cleavage were assessed in mixed supernatants and lysates by western blot. **C** Fixed-Airyscan confocal imaging of WT BMDM primed for 4 hours with 1 µg/mL Pam3CSK4, and incubated with HEPES (Ctrl) or 10 µM of TopFluor CL (BODIPY-CL) for a further 18 hours. Macrophages were immunostained with TOMM20 (red), BODIPY-CL (green) and DAPI (grey). Images are maximum intensity projections of Z-stack acquisitions. Scale bar= 10 µm. **D** WT BMDM were incubated for 4 hours with 1 µg/mL Pam3CSK4. Cells were then lysed and incubated with HEPES (-), 100 or 200 µg/mL CL or 100 µg/mL LPS for 1 hour. 1 µg/mL of biotinylated LPS (biotin- LPS), was added, and lysates were incubated for further 2 hours. Biotinylated LPS with bound proteins was purified using magnetic streptavidin beads. Both purified (bound) and unbound fractions were immunoblotted for CASP11 and GAPDH by western blot. **E - F Upper panels** Schematic of full-length (E) and ΔCARD (F) CASP4 constructs with the DmrB dimerisation system, permitting controlled dimerisation by AP20187 (AP). All DmrB constructs were N- terminally V5-tagged. **Lower panels** HEK293T cells were transfected with the native or the catalytic cysteine mutant (C258A) form of V5-DmrB-CASP4 (full length, E) or V5-DmrB-τιCARD-CASP4 (F) constructs. 24 hours later, cells were harvested, plated, and 4 hours later treated with AP for 30 min in the absence or presence of 10 µM CL. CASP4 auto-processing was analysed by western blot of cell lysates. **G** Recombinant CASP4-CARD-eGFP (1 µM) was incubated for 1 hour at 37°C with HEPES (Ctrl), or 1.67 mM of liposomes containing phosphatidylcholine (PC) or CL alone or increasing CL:PC molar ratio (5:95, 25:75, 50:50 and 75:25) (total lipid concentration). Samples were centrifuged and the GFP fluorescence was measured in supernatant (unbound CARD in non-liposome fraction) and resuspended pellet (liposome-bound CARD). The Y-axis represents the percentage of overall fluorescence in the pellet. **Data information:** Each symbol is the mean of technical triplicates from an independent biological replicate. Bars are the mean of three independent biological replicates (n = 3) ± SEM . Blots and images are representative of three or four independent experiments (n = 3 to 4). **Statistical analysis: G** One-way ANOVA Dunnett’s multiple comparisons test. Significant difference for p<0.05 (*), not significant p≥0.05.

Pam-priming did not affect CASP4 expression in HMDM but upregulated CASP11 in BMDM, and pro-IL-1β in both HMDM and BMDM (Fig 2 A-B), confirming previous findings at the mRNA level (Schroder et al., 2012). In BMDM, pro-IL-1β expression was further upregulated by LPS, likely by TLR4 activation (Fig 2 B). To confirm that CL did not suppress CASP4/11 signalling via an indirect mechanism involving TLR4, we measured TNF secretion (Fig S2 A-B) alongside inflammasome signalling outputs (Fig 2 A-B). CL did not affect LPS-induced TNF secretion in HMDM and BMDM (Fig S2 A-B), in line with our previous report that CL does not inhibit TLR4 signalling induced by the LPS dose used here to activate CASP4/11 (Pizzuto et al., 2019). Further, in Pam-primed TLR4-deficient BMDM (*Tlr4*^-/-^), CL suppressed iLPS-induced signalling outputs, including cleavage of CASP11, GSDMD, CASP1, and IL-1β (Fig S2 C) as well as LDH and IL-1β release (Fig S2 D-E). Thus, CL inhibits CASP11 signalling independently of TLR4.

The efficacy of LPS-induced CASP4/11 activation depends on the amount of LPS delivered into the cytosol (Hagar et al., 2013; Kayagaki et al., 2013). To address the possibility that CL interfered with LPS delivery to the cytosol by transfection agents or CTB, we used alternative means to deliver LPS intracellularly using FuGENE (Fu), Xfect or electroporation. CL suppressed iLPS-induced LDH and IL-1β release in primed BMDM (Fig S3 A-F), confirming that CL inhibits CASP4/11 signalling regardless of the intracellular LPS delivery system. To evaluate the contribution of CASP11 in response to electroporated LPS, we primed and electroporated *Casp11*^-/-^ BMDM in parallel. Electroporation of WT BMDM in the presence of LPS resulted in a significant increase in LDH and IL- 1β release, which was reduced by CL to the level measured in control and *Casp11*^-/-^ electroporated cells (Fig S3 E-F). Thus, CL suppresses iLPS-induced CASP11 signalling outputs even when LPS carriers are not used, further validating CL as a *bona fide* CASP11 inhibitor.

To determine whether exogenously added CL relocates to the cytosol, we cultured BMDM with a modified form of CL that is covalently attached to a fluorescent probe, and found that CL was internalised by macrophages (Fig S3 G). A previous study showed that exogenous CL localises to mitochondria using the mitochondrial dye MitoTracker (Ikon et al., 2015), which we confirmed by showing that exogenous CL co-localises with TOMM20 in the mitochondrial outer membrane (Fig 2 C). These data indicate that CL is taken up by cells and accumulates in the mitochondrial outer membrane, from which it has access to cytosolic proteins such as CASP4/11.

In the context of a bacterial infection, guanylate-binding proteins (GBP) promote LPS binding to CASP4/11 (Kirkby et al., 2023; Santos et al., 2020; Tretina et al., 2019). We sought to determine whether CL could inhibit LPS-induced noncanonical inflammasome indirectly, by targeting GBPs. Thus, we tested BMDM deficient in the chromosome 3 GBP (*Gbp*^chr3-/-^), in which the cluster of GBPs that promote CASP11 responses to bacteria is deleted (*Gbp1, Gbp2, Gbp3, Gbp5, Gbp7,* and *Gbp2ps)* (Enosi Tuipulotu et al., 2023; Meunier et al., 2014; Santos and Broz, 2018; Yamamoto et al., 2012). We treated Pam-primed *Gbp*^chr3-/-^ BMDM with iLPS in the absence or presence of CL. CL suppressed, iLPS-induced IL-1β, LDH and IL-18 release in *Gbp*^chr3-/-^ BMDM (Fig S3 H-J). These data demonstrate that these GBPs are dispensable for the suppressive effect of CL on noncanonical inflammasome signalling.

Given that exogenous CL co-localises with the mitochondrial outer membrane, and the inhibitory activities of CL do not involve GBPs, we hypothesised that, once in the cytosol, CL may prevent CASP4/11 activation by blocking LPS interaction with CASP4/11. To test this, we incubated the cytosolic fraction of BMDM with biotinylated LPS in the absence or presence of CL (or unconjugated LPS as a control), and then pulled down biotinylated LPS using streptavidin beads. LPS pulled down CASP11 in CL-untreated cells, and this was suppressed by CL in a dose-dependent manner, similar to that of unconjugated LPS (Fig 2 D). Thus, CL prevents LPS binding to CASP11, thereby suppressing LPS-induced CASP11 activation. This led us to hypothesise that CL may block the interaction between LPS and CASP4/11 by targeting either of these interaction partners. To test whether CL target CASP4, we employed the DmrB dimerisation system that enables drug-inducible, LPS-independent CASP4 activation. Here, we transfected HEK293T cells with a construct encoding full-length CASP4, N-terminally fused with a V5-tagged DmrB domain (V5-DmrB-FL_CASP4). Cells were treated with the dimeriser drug AP20187 (AP) to induce DmrB-mediated dimerisation, autocleavage and activation of CASP4. Cells were treated with AP in the presence or absence of CL, and cell lysates were assessed for CASP4 cleavage. CL inhibited AP-induced CASP4 cleavage (Fig 2 E), indicating that CL indeed suppresses CASP4 signalling independently of LPS.

Our data indicating that CL inhibits CASP4 independently of LPS while preventing LPS binding to CASP11 suggest that CL competes with LPS for binding to CASP4/11. Given that LPS interacts with CARD domain of CASP4/11 (Shi et al., 2014), we reasoned that CL may target the CARD domain to block LPS interactions and resultant CASP4/11 activation. To test this, we expressed a V5- DmrB-tagged variant of CASP4 that lacks the CARD domain (V5-DmrB-DCARD- CASP4) in HEK293T cells and induced CASP4 activation with AP, in the absence or presence of CL. When CASP4 lacked its CARD domain, CL failed to suppress AP-induced CASP4 cleavage (Fig. 2 F). For both full-length and DCARD CASP4 constructs, we verified that AP-induced CASP4 cleavage represented CASP4 auto-processing activity, as this cleavage was blocked when we mutated the catalytic cysteine (C258A) (Fig. 2 E-F). Thus, CL targets the CASP4 CARD domain rather than the protease domain to suppress noncanonical inflammasome signalling.

To determine whether CL directly binds to the CASP4 CARD, we employed a non-cellular, fully-recombinant system. We generated recombinant protein for the CARD domain of CASP4 tagged with eGFP (CARD-eGFP), and incubated this with liposomes containing phosphatidylcholine (PC or CL alone, or varying ratios ratio of CL to PC) and determined whether the CASP4 CARD interacts specifically with CL. High-speed centrifugation then separated liposomes from unbound CARD-eGFP. CARD-eGFP levels were quantified by GFP fluorescence in both liposomal (bound) and soluble (unbound) fractions. While liposomes containing PC alone showed no fluorescence, liposomes containing 75 and 100% CL exhibited significant fluorescence (Fig 2 G), indicating that CL binds to the eGFP-tagged CASP4 CARD. In all, these data demonstrate that CL binds the CASP4 CARD domain.

### Cardiolipin specifically inhibits CASP4/11 and not CASP1

Like CASP4/11, CASP1 also contains a CARD that is required for its activation. To test whether CL inhibits CASP1 activation, we treated primed BMDM with NLRP3/CASP1 activators (nigericin, silica) or the AIM2/CASP1 activator poly(dA:dT), in the absence or presence of CL. CL did not affect IL-1β and LDH release induced by nigericin, silica or poly(dA:dT) in murine macrophages (Fig 3 A-B), or nigericin-induced IL-1β release in HMDM (Fig 3 C). Thus, CL specifically inhibits CASP4/11 without affecting CASP1, NLRP3 or AIM2 activity in macrophages.

**Figure 3.**
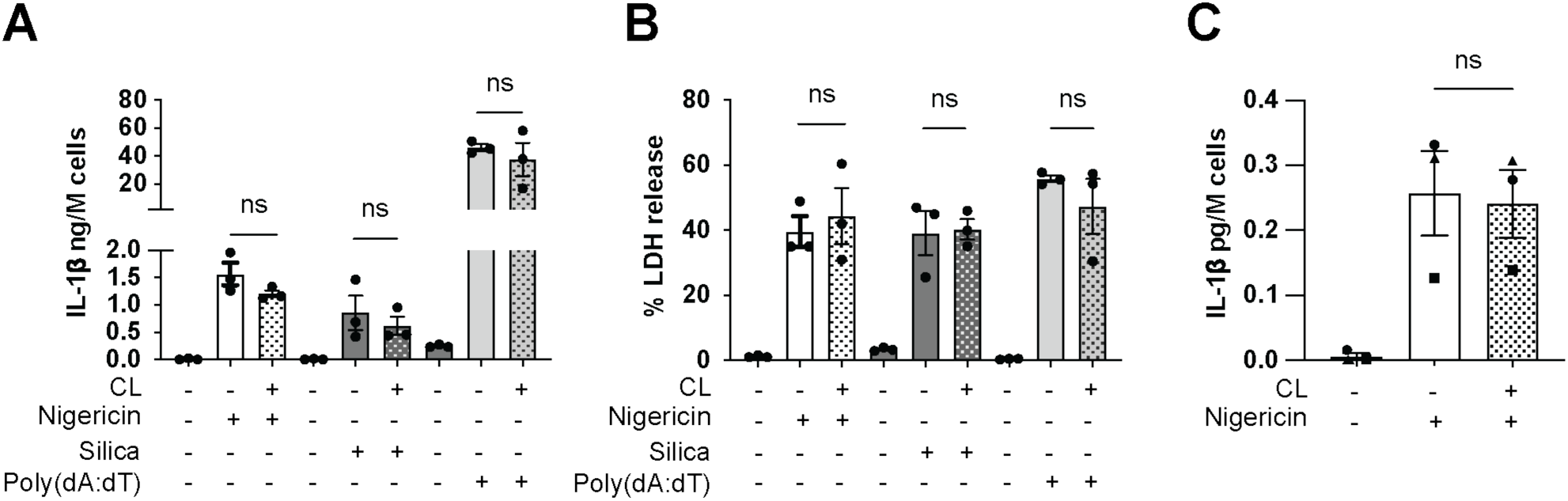
Cardiolipin inhibits CASP4/11 but not CASP1 activation. WT BMDM (**A-B)** and HMDM (**C**) were primed for 4 hours with 1 µg/mL Pam3CSK4. Cell culture medium was then replaced with 10 µM nigericin, 200 µg/mL silica or 5 μg/ml of poly(dA:dT) in the presence of 10 μM CL (+) or HEPES vehicle (-). Cells were incubated for 1 hour (nigericin), 18 (silica), or 4 (poly(dA:dT)) hours. Cell supernatants were analysed for cleaved IL-1β (ELISA) and LDH (cytotoxicity assay). **Data information:** Each symbol is the mean of technical triplicates from an independent biological replicate. Bars are the mean of three independent biological replicates (n = 3) ± SEM. **Statistical analysis:** Unpaired t-test. Statistical significance was defined as follows: p ≥ 0.05 not significant (ns).

Given the absence of CASP4/11-specific inhibitors and the wide variety of specific NLRP3 inhibitors (Coll et al., 2022), inflammasome-targeting research and clinical trials focus on the NLRP3-CASP1 axis rather than CASP4/11 (Coll et al., 2022; Hagar et al., 2013; Kayagaki et al., 2011; Schmid-Burgk et al., 2015). Our discovery that CL is a specific CASP4/11 inhibitor offers a tool to allow CASP4/11 functions to be studied in diverse inflammatory pathways and diseases, including sepsis. Such specificity may be crucial, as generic IL-1β pathway inhibition in clinical trials for several diseases increased the risk of opportunistic infections (Lopalco et al., 2016; Marshall, 2014; Salliot et al., 2009; Winthrop, 2006). Inhibitors that block the CASP4/11 active site are poorly selective due to similarities within the active site between inflammatory caspases (Ekert et al., 1999; Green, 2022). Here, we show that CL is an inhibitor of CASP4/11, that targets the CARD domain of CASP4 and likely CASP11 to prevent CASP4/11, but not CASP1, activation. This discovery gives proof-of-concept that targeting the CARD can allow specific inhibition of CASP4/11, and perhaps other inflammatory caspases. Indeed, the CASP4/11 CARD domain differs substantially from CASP1, as the CASP4/11 CARD binds LPS while the CASP1 CARD does not (Devant et al., 2021; Shi et al., 2014), and CARD recruitment to ASC filaments is required for CASP1 but not CASP11 activation (Ross et al., 2022).

### Cardiolipin mitigates endotoxemia-induced pro-inflammatory cytokine secretion *in vivo*

We next sought to determine whether CL blocks pathological noncanonical inflammasome responses *in vivo*, using a murine endotoxemia model that engages CASP11 signalling (Kayagaki et al., 2013; Napier et al., 2016). We intraperitoneally administered CL (25 µg/g, versus vehicle) immediately prior to LPS (10 µg/g) challenge in wild-type and *Casp11*^-/-^ mice. LPS-induced inflammatory responses (TNF, IL-1β in sera) and weight loss were measured 6 hours post-injection. TNF and IL-1β in serum and changes in weight loss were undetectable in sham (PBS)-challenged animals regardless of CL administration, and significantly increased by LPS challenge (Fig 4). LPS-induced serum TNF levels were not dependent on CASP11 and were not suppressed by CL (Fig 4A), in line with our earlier findings that CL does not affect TLR4 activation induced by high LPS concentrations ((Fig. S2 A-B) and (Pizzuto et al., 2019)). In a murine model of polymicrobial sepsis, mice lacking both TLR4 and CASP11 showed higher mortality than those only lacking CASP11 (Deng et al., 2018). This suggests that preserving TLR4-dependent anti-bacterial defence while blocking CASP4/11-induced cell death and IL-1β production could offer benefit for treating sepsis. LPS-induced levels of circulating IL-1β were significantly reduced in *Casp11*^-/-^ compared to wild-type mice, demonstrating the engagement of CASP11 noncanonical inflammasome by LPS (Fig 4B). CL significantly suppressed LPS-induced IL-1β production to levels similar to *Casp11*^-/-^ mice (Fig 4B). Thus, CL suppresses noncanonical inflammasome signalling during murine endotoxemia. This discovery opens new avenues towards blocking LPS/CASP4/11 signalling sequelae such as organ damage and lethality (Chen et al., 2019; Cheng et al., 2017; Deng et al., 2018; Hagar et al., 2013; Kajiwara et al., 2014; Kayagaki et al., 2015, 2011; Tang et al., 2018; Wei et al., 2024).

**Figure 4:**
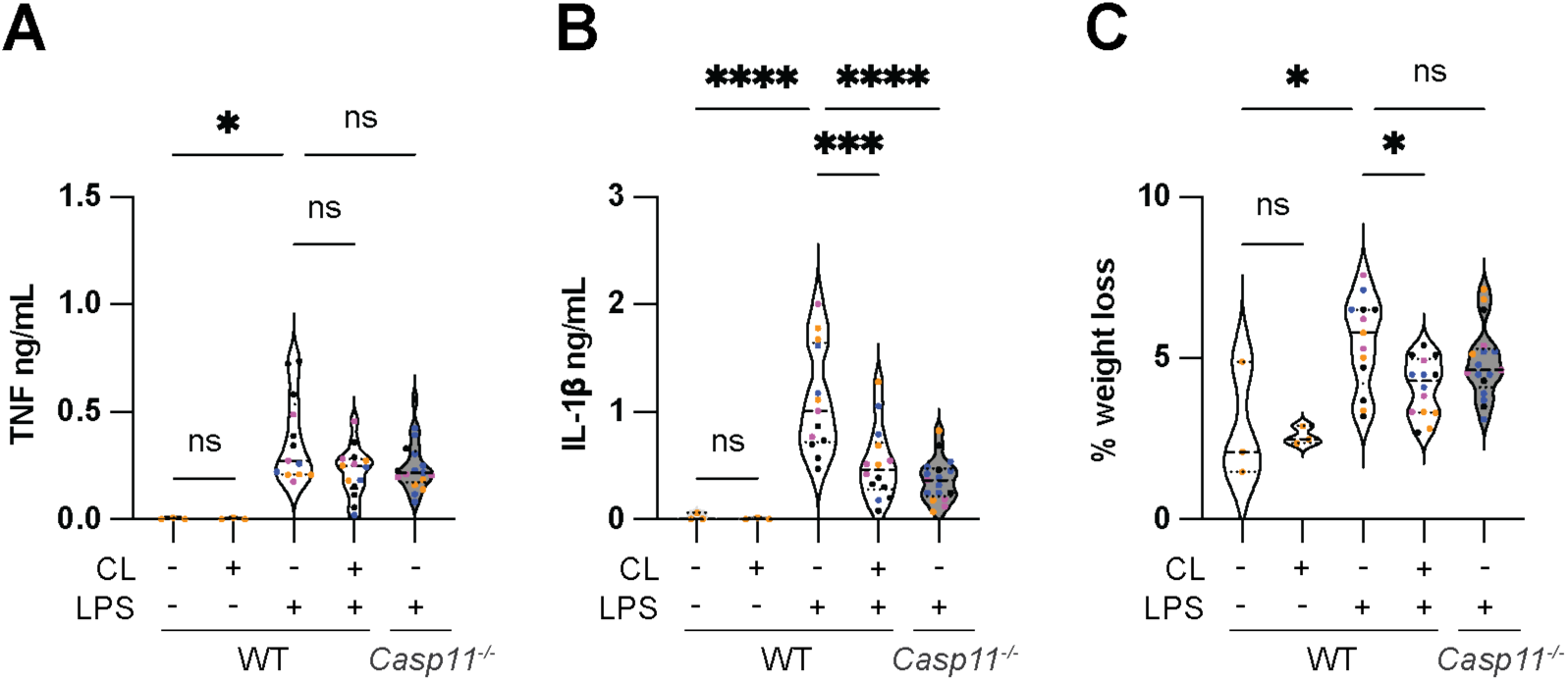
Cardiolipin mitigates endotoxemia-induced systemic inflammation *in vivo*. WT and *Casp11^-/-^* mice were weighed and injected (i.p.) with HEPES or 25 µg/g CL. 10 minutes later, mice were i.p. challenged with PBS or 10 µg/g LPS. After 6 hours, the animal’s weight was recorded again, and blood was collected. TNF (A) and IL-1β (B) were quantified in sera by ELISA. Body weight loss was calculated as the percentage of the initial weight (C). **Data information:** Violin plot of data from four different cohorts of mice (individual mice from each cohort shown as colour-matched data points). WT PBS and WT PBS + CL n=3; WT HEPES + LPS n=13; WT CL + LPS n=13; and *Casp11^-/-^*HEPES + LPS n = 17. **Statistical analysis: A and C** One-way ANOVA Sidak’s multiple comparisons test. **B** Krustal-Wallis with Dunn’s multiple comparisons test. Statistical significance was defined as follows: 0.05 > p > 0.001 (*), 0.001 ≤ p > 0.0001 (**), p ≤ 0.0001 (***), p ≥ 0.05 not significant (ns).

CL also significantly blunted LPS-induced weight loss (Fig 4 C), supporting the safety of CL when injected i.p. in mice, and highlighting its protective effects in suppressing inflammasome-driven cytokine production and endotoxemia-induced weight loss (Fig. 4 BC). CL is naturally present in the body as a key mitochondrial lipid, which may contribute to its favourable safety profile (Oemer et al., 2018). Safety and efficacy make CL an attractive candidate for developing therapies to inhibit LPS/CASP4/11 signalling sequelae such as organ damage and lethality (Chen et al., 2019; Cheng et al., 2017; Deng et al., 2018; Hagar et al., 2013; Kajiwara et al., 2014; Kayagaki et al., 2015, 2011; Tang et al., 2018; Wei et al., 2024). When considering CL as a therapeutic, the route of administration requires careful consideration. Intra-tracheal administration of CL following LPS exposure in a pneumonia murine model resulted in CL degradation which produced pro-inflammatory CL metabolites, and lung injury (Chakraborty et al., 2017; Ray et al., 2010). By contrast, intraperitoneal (i.p.) administration of CL showed no adverse effects, by us and others (Ikon et al., 2018; Ordóñez- Gutiérrez et al., 2015), demonstrating safe *in vivo* application via this route.

In summary, this study shows that natural CL, unlike RS-LPS, is a specific inhibitor of CASP4/11 with *in vivo* efficacy that may be exploited in new approaches for suppressing endotoxemia-induced organ damage and lethality. Here, one key advantage of CL is that it does not block TLR4 or canonical inflammasome responses, and is thereby unlikely to compromise these key pathways of antimicrobial defence. By identifying CL as a specific CASP4/11 inhibitor, we provide a tool reagent to study the involvement of CASP4/11 in inflammatory pathways both *in vitro* and *in vivo*. Ultimately, the discovery that CL inhibits CASP4/11 may have important implications in future studies of noncanonical inflammasome regulation by bacteria and mitochondria. Such future studies may provide the missing links to explain why modifications of endogenous CL are associated with inflammatory disorders in diseases such as Barth Syndrome (Pizzuto and Pelegrin, 2020).

## Materials and Methods

### Cardiolipin liposome preparation

The cardiolipin used in this work is double unsaturated cardiolipin (18:2) extracted from bovine heart, purchased from Avanti Polar Lipid (840012P) or Merck (C0563). CL was dissolved in CHCl3 (Merck, 650498) at 1 mg/mL, and lipid films were prepared by solvent evaporation under a filtered nitrogen stream before being dried overnight and stored at −20°C. Before each experiment, liposomes were freshly formed by resuspending lipid films with filtered 10 mM HEPES (Gibco, 15630080), heating for 20 minutes at 70°C, and then sonicating for 5 min at 44 kHz in an ultrasonicator bath (Grant XUBA1).

### Intracellular delivery of LPS or poly(dA:dT)

Poly(dA:dT) (InvivoGen, tlrl-patn-1) or ultrapure LPS from *Escherichia coli* 0111:B4 (InvivoGen, tlrl-3pelps or ALX-581-014-L002, Enzo Life Science), *Escherichia coli* K12 (InvivoGen, tlrl-peklps), *Salmonella enterica serovar minnesota mutant* R595 (InvivoGen, tlrl-smlps), *Pseudomonas aeruginosa* (Merck L8643), or *Rhodobacter spheroides* (InvivoGen, tlrl-prslps) were first mixed with FuGENE ®HD (Promega, E2311), Lipofectamine™ LTX Reagent with PLUS™ Reagent (LTX) (Invitrogen, A12621), Xfect (Clontech Laboratories, 631318), or cholera toxin B (CTB) (Merck, C9903) in a small volume (1/10^th^ final volume) of Opti-modified Eagle’s medium (MEM)™ reduced-serum medium (OptiMEM) (Invitrogen, 31985-070,) that was previously heated to 37°C. The mix was vortexed, and then incubated for 15 minutes at room temperature to allow complexes to form, and then added to cells in OptiMEM™. Final concentrations are indicated in the figure legends.

### HEK293T cell culture, transfection and treatments

All cells were cultured in humidified incubators at 37°C and with 5% CO2. HEK293T (ATCC CRL-3216) in Dulbecco’s modified Eagle’s medium (DMEM; Gibco) supplemented with 10% heat-inactivated fetal bovine serum (FBS) and 1% penicillin-streptomycin (Pen-Strep).

The full-length coding sequence of human *CASP4* (residues 1-377) and delta- CARD *CASP4* (residues 81-377) were cloned as N-terminal DmrB fusions into the mammalian pEF6 expression vector (Invitrogen), as either the wild-type sequence or with an inactivating mutation in the catalytic cysteine (C258A). Sequences were cloned in-frame with an N-terminal V5 tag and a C-terminal HA tag.

The DmrB constructs were transfected into HEK293T cells seeded in 10 cm cell culture dishes using lipofectamine 2000 (Thermo Fisher). After overnight expression, the cells were harvested and re-seeded at 0.3 x10^6^ cells per well in 24 well plates in OptiMEM. After 3 hours, 30 µM CL was added (or HEPES vehicle), and 1 hour later OptiMEM or the dimeriser drug AP20187 (AP, 1 µM) was added. 30 minutes later, cell lysates were collected, and CASP4 expression and cleavage were assessed by western blot using the V5 antibody.

### HBEC cell line culture and treatments

Immortalised human bronchial epithelial cells HBEC-KT (ATCC CRL-4051) were maintained in complete keratinocyte medium (Gibco 17005042, supplemented with the provided bovine pituitary enzyme and epidermal growth factor) and passaged at 70-80 % confluency. Cells were seeded at 0.1 x10^6^ cells in 100 μL per well in 96 well plates in complete keratinocyte medium supplemented with 1 μg/mL Pam3CSK4. After 18 hours, the cell culture medium was replaced with OptiMEM alone or containing 10 µM of the NLRP3 inhibitor MCC950 or the CASP1/4 inhibitor VX765 (MedChem Express). 1 hour later, OptiMEM or 5 μg/mL of LPS complexed with 0.25% LTX were added in the absence or presence of 10 µM CL . 4 hours later, supernatants were collected, centrifuged at 600 g, and assayed for cytokine and LDH release.

### Differentiation of bone marrow-derived macrophages (BMDM)

Experiments conducted at the Biomedical Research Institute of Murcia used wild-type or *Nlrp3^−/−^* C57BL/6J male and female mice between 8 and 13 weeks of age and bred under specific pathogen-free (SPF) conditions, in accordance with the Hospital Clínico Universitario Virgen Arrixaca animal experimentation guidelines, and the Spanish national (RD 1201/2005 and Law 32/2007) and EU (86/609/EEC and 2010/63/EU) legislation. Accordingly, no specific procedure approval is needed when animals are sacrificed to obtain biological material. Mice were euthanised by CO2 inhalation and the bone marrow was flushed from the leg bone cavity and resuspended in differentiation medium (DMEM medium, with L-glutamine, without sodium pyruvate (Biowest, 11320) supplemented with 10% heat-inactivated fetal bovine serum premium (Biowest, A5256701), 2 mM glutamine (Lonza, BEBP17-605E), 50 U/mL penicillin, 50 μg/mL streptomycin (PEN-STREP, Lonza, 17-603 DE17-603), and 20% of supernatant from L929 cultures. The bone marrow cell suspension was maintained in Petri dishes in a 37°C/5% CO2 atmosphere. After 2 days, the differentiation medium was supplemented, and cells were maintained for four extra days, before replating in DMEM medium, with L-glutamine, without sodium pyruvate supplemented with 10% heat-inactivated fetal bovine serum premium (Biowest, A5256701).

Alternatively, experiments at the Institute for Molecular Bioscience (University of Queensland) used wild-type, *Tlr4^−/−^, Casp11^−/−^* and *Nlrp3*^−/−^ C57BL/6J male and female mice between 6 and 14 weeks of age and bred under specific pathogen-free (SPF) facilities at the University of Queensland. All protocols involving mice were approved by the University of Queensland Animal Ethics Committee and compliance with relevant ethical regulations was ensured (2023/AE000019, 2023/AE000020, 2020/AE000419). Mice were euthanised by CO2 inhalation, the bone marrow was flushed from the bone cavity, filtered, centrifuged at 400 g for 5 minutes and resuspended in differentiation medium consisting of RPMI 1640 medium (Life Technologies, 11875093) supplemented with 10% heat-inactivated and endotoxin-free fetal calf serum (FCS) (Gibco), 2 mM GlutaMAX (Life Technologies, 35050061), 50 U per ml penicillin–streptomycin (Life Technologies, 15140122) and 150 ng/ml recombinant human macrophage colony-stimulating factor (CSF-1; endotoxin-free, expressed and purified by the University of Queensland Protein Expression Facility). . After five days, the differentiation medium was supplemented, and cells were maintained for a further day before replating for experiments in cell differentiation medium.

Experiments conducted at the Australian National University (Xfect/LPS/polydAdT transfection) used C57BL/6NCrlAnu mice and *Gbp^chr3-/-^*mice (Yamamoto et al., 2012) sourced from Osaka University. Primary BMDM were differentiated and cultured in Dulbecco’s Modified Eagle Medium (DMEM) (11995073, Gibco Thermo Fisher Scientific) with 30% L929-conditioned medium 1% penicillin and streptomycin (10378016, Gibco Thermo Fisher Scientific) and 10% foetal bovine serum (FBS; F8192, Sigma).

### BMDM stimulation

BMDM differentiated for six days were washed and harvested using Dulbecco’s phosphate-buffered saline (PBS) (Thermo Fisher Scientific, J67670.K2). The cells were then counted, centrifuged for 5 minutes at 500 g and resuspended in full medium to a concentration of 1 × 10^6^ cells/mL and distributed in 96-well plates (100 μL/well), 24-well plates (500 μL/well), or 6-well plates (2 mL/well) and cultured overnight. Medium was added, either alone or supplemented with 1 μg/mL Pam3CSK4 (InvivoGen, tlrl-pms). After 4 hours, supernatants were collected, and stimulants diluted in OptiMEM were added as indicated in a final volume of 100 μL/well (96-well plates), 250 or 300 μL/well (24-well plates) or 800 μL/well (6-well plates). After treatment, the supernatants and cell lysates were collected.

### Macrophage CASP11-LPS binding assay

BMDM were primed with 1 μg/mL of Pam3CSK4, then lysed in 100 μL lysis buffer (50 mM HEPES pH 7.4, 150 mM NaCl, 2 mM EDTA, 10% glycerol, and 1% Triton X-100, plus protease inhibitor 100 μL/mL) per million cells. Lysates were spun at 13,000 g for 10 minutes, and the pellet containing debris and non-lysed cells was discarded. The lysate was aliquoted (250 μL per tube), and reagents added to the same volume: CL (0, 100 or 200 μg), or LPS (0 or 100 μg). Tubes were incubated at room temperature for 2 hours on a Benchmark rotating wheel, and then biotinylated LPS (0 or 1 μg/mL) was added. 1 hour later, 1% vol/vol of Promega Z5481 streptavidin magnetic beads were added. Beads were pre-washed with BSA-T buffer (0.15 % tween, 5% Bovine Serum Albumin in PBS) and blocked by incubation for one hour in BSA-T buffer on a rotating wheel. Samples were incubated with beads overnight at 4 °C. Then, supernatants (unbound fractions) were separated from beads. Beads were recovered and washed four times for 5 minutes in BSA-T buffer at room temperature on the rotating wheel. Then beads were recovered, and bound samples were eluted from beads by adding 40 μL of lysis buffer diluted 3:4 in NuPAGE 4X (final NuPAGE concentration 1X) and boiling at 100 °C for 5 minutes. Beads were recovered from preservative solution, buffer or samples by gentle magnetic separation using a DynaMag-2 magnetic rack (Invitrogen). Unbound fractions were diluted 3:4 in NuPAGE 4X (final NuPAGE concentration 1X) and boiled at 100 °C for 5 minutes. The amount of CASP11 in LPS-bound and unbound fractions was assessed by Western Blot.

### Differentiation of human macrophages from human monocytes (HMDM)

Studies using primary human cells were approved by the UQ Human Research Ethics Committee (HE000413). The Australian Red Cross Blood Service provided buffy coats from anonymous, informed and consenting adults for this research study. Human monocytes were isolated from screened buffy coats by density centrifugation with Ficoll-Plaque Plus (GE Healthcare) followed by Miltenyi Biotec magnetic-assisted CD14+ cell sorting, according to manufacturer protocols. Monocytes were differentiated to macrophages by 6 days culture at 37 °C with 5% CO2 in Iscove’s Modified Dulbecco’s Medium (IMDM; Gibco) medium supplemented with 10% endotoxin-free heat-inactivated fetal bovine serum (Gibco), 1% penicillin-streptomycin, 1x GlutaMAX supplemented with recombinant human CSF-1 (150 ng/mL; endotoxin-free, produced in insect cells by the UQ Protein Expression Facility).

### HMDM stimulations

HMDM differentiated for 6 days were washed and detached from their dishes using Dulbecco’s phosphate-buffered saline (PBS) (Thermo Fisher Scientific, J67670.K2). The cells were then counted, centrifuged for 5 minutes at 500 g, and resuspended in full medium to a concentration of 0.5 × 10^6^ cells/mL and distributed in 96-well plates (100 μL/well) or 12-well plates (1 mL/well) and cultured overnight . Medium was added, either alone or supplemented with 1 μg/mL Pam3CSK4 (InvivoGen, tlrl-pms). After 4 hours, supernatants were collected, and stimulants diluted in OptiMEM were added as indicated in a final volume of 100 μL/well (96-well plates) or 800 μL/well (12-well plates). After treatment, the supernatants and cell lysates were collected.

### Cytokine assays

Murine IL-18 was quantified with the ELISA™ Kit (Thermo Fisher Scientific, BMS618-3TEN). Murine TNF and IL-1β were quantified in cell supernatants using ELISA (DuoSet R&D Systems: DY401, D410 or Invitrogen: 88-7324-88, 88- 7013A-88). Human TNF, IL-1β, and IL-18 were quantified in cell supernatants using the DuoSet ELISA Kit from R&D Systems (DY210, DY201 and DY318).

Absorbance was read with a Tecan Microplate Reader or a BioTek Synergy HT Microplate Reader. For *in vitro* experiments, cytokine amounts were reported as ng or pg per million cells to standardise the difference in cell amount/volume of media ratio between the different plate layouts.

### LDH assay

LDH activity was quantified in cell supernatants using the Cytotoxicity Detection Kit (Merck), following the manufacturer’s instructions. Absorbances at 492 and 620 nm were measured with a BioTek Synergy HT Microplate Reader every minute for 20 minutes, with the slopes of increase in absorbance calculated with respect to time, and background values subtracted from the value of each supernatant reported as percentages of the sum of the value measured in the supernatant and the lysate of the untreated condition (total LDH). Untreated cell lysates were also obtained to estimate total cellular LDH, and were prepared as follows: cells were lysed with 2% Triton lysis buffer comprising 150 mM NaCl (Sigma-Aldrich), 2% Triton X-100 (Sigma-Aldrich), and 50 mM Tris-HCl pH8 (Thermo Fisher Scientific) supplemented with 100 µL/mL of protease inhibitor (Sigma-Aldrich). Cells were harvested by scraping in cold lysis buffer on ice. Lysates were then incubated for 30 minutes on ice with a vortex every 10 minutes, before centrifugation for 10 minutes at 13,000 g in a microcentrifuge (1-14K, Sigma) to remove cell debris.

Alternatively, LDH activity was quantified in all cell supernatants using the Cytox96 non-radioactive cytotoxicity assay (Promega). Absorbances at 492 and 620 nm were measured with a Tecan Microplate Reader and reported as percentages of the value measured in the supernatant of cells treated with 0.1% triton for 10 minutes (total LDH).

### Western Blotting

Cells were lysed in complete lysis buffer (66 mM Tris-Cl pH 7.4, 2% SDS, 100 mM DTT, Benzonase 0.01% vol/vol, NuPAGE 1X). Cell supernatants were concentrated following CHCl3/MeOH precipitation as described earlier (Groß, 2012) and resuspended in complete lysis buffer. Samples were incubated for 5 minutes at 100°C, then resolved by SDS–PAGE using 4–20% Mini-PROTEAN TGX stain-free gels (BioRad) and transferred onto nitrocellulose membrane using the Trans-Blot Turbo transfer system (BioRad). Membranes were blocked in 5% (wt/vol) dried milk in TBS-T (10 mM Tris/HCl, pH 8, 150 mM NaCl, 0.05% vol/vol Tween-20) for 1 hour at room temperature. Membranes were incubated for 18 hours at 4°C with primary antibody diluted in 5% (wt/vol) dried milk in TBS-T and then 1 hour at room temperature with the appropriate horseradish peroxidase (HRP)-conjugated secondary antibody diluted in 5% (wt/vol) dried milk in TBS-T for 1 hour. Membranes were developed using SuperSignal West Femto Maximum Sensitivity Substrate ultra-sensitive enhanced chemiluminescent (ECL) (Thermo Scientific). Membranes were then visualised using a ChemiDoc MP Imaging System with Image Lab 6.1 (BioRad). Secondary antibodies on membranes were inactivated by incubation with 30% hydrogen peroxide for 20 minutes before re-probing. The following primary antibodies were used at 1:1000: anti-IL1beta (RnD AF-401-NA), anti-CASP1p20 (AG-20B-0042, Adipogen), anti-GSDMD (Ab209845, Abcam), anti-V5-tag Sv5-PK1 (ab27671, Abcam), anti-CASP11 (ab180673, Abcam), and anti-CASP4 (56056, Santa Cruz). Secondary HRP-conjugated antibodies used were anti-rabbit IgG or anti-mouse IgG (7074S, 7076S; both 1:5,000 Cell Signaling Technology) or anti-goat IgG at 1:10,000 (Abacus). Tubulin and GAPDH were blotted using Rhodamine-conjugated anti-tubulin and anti-GAPDH (BioRad) at 1:10,000 protected from light.

### Expression and isolation of recombinant CASP4 CARD

The CASP4 CARD domain (1-80 aa), codon-optimized for *E. coli*, was expressed with C-terminal eGFP-6xHis tag from pET28b vector in ClearColi BL21(DE3) (Lubio Science), grown in Luria-Bertani medium. Expression was induced at OD600 of 0.5 with 0.2 mM isopropyl-b-D-galactopyranoside (IPTG) at 18°C, overnight. For purification, all glassware was first rinsed with 1M NaOH, to avoid contamination by lipopolysaccharide. Harvested bacteria were resuspended in IMAC-A buffer (20 mM Tris-HCl pH7.9, 300 mM NaCl, 20 mM imidazole, 5 mM B-mercaptoethanol, 1% Tween-20) and lysed by sonication. The CASP4-CARD- GFP-His was purified by IMAC affinity chromatography on Ni Sepharose^®^ 6Fast Flow (Cytiva), eluted using 300 mM imidazole in IMAC-B buffer (20 mM Tris-HCl pH7.9, 300 mM NaCl, 300 mM imidazole). Finally, monomeric CASP4-CARD- eGFP-His was further purified to homogeneity by size-exclusion chromatography, using Superdex-75 10/300 equilibrated in SEC buffer (20 mM HEPES-NaOH pH7.5, 300 mM NaCl, 10% glycerol, 2 mM dithiothreitol) and frozen in liquid nitrogen.

### Liposome co-sedimentation assay for CASP4-CARD binding assay

To analyse the binding of CASP4-CARD to CL, 1 µM recombinant CARD in binding buffer (20 mM HEPES-KOH pH 7.5, 150 mM KCl) was mixed with liposomes containing 1.67 mM total lipid, with increasing proportions of CL (0 to 100 % mol/mol) and decreasing proportions of PC (L-alpha-phosphatidylcholine, egg, chicken, Avanti Lipids), and incubated at 37°C for 1 hour with constant shaking. Subsequently, mixtures were centrifuged at 20,000 g at 4°C for 30 minutes. Liposome-containing pellets were resuspended in binding buffer, using a volume equal to the supernatant volume. The GFP fluorescence (ex: 485 nm/ em: 528 nm) was measured, using the microplate reader Cytation5 (Biotek).

### Immunofluorescence microscopy

BMDM were plated on 1’ glass coverslips at 2 x 10^5^ cells/ml and fixed in 4% PFA (Pierce) for 15 minutes at 37°C. Cells were permeabilised in 0.1% saponin (Sigma Aldrich) and non-specific binding was blocked using 0.5% BSA (Sigma Aldrich) for 30 minutes, followed by incubation with primary and secondary antibodies for 1 hour. The primary antibody used was anti-TOMM20 (ab186735, Abcam; 1:400), together with the actin probe phalloidin-594 (6.6µM, Invitrogen) and DAPI (0.1 µg/ml, Sigma Aldrich). Alexa Fluor-594 (A21442) was purchased from Molecular Probes. Coverslips were mounted with ProLong Gold Antifade Reagent (Invitrogen). Images were acquired on a Zeiss Axiovert 200 Inverted Microscope Stand with LSM880 Confocal Scanner running Zeiss Zen 2012 Black software. The microscope was equipped with 405, argon ion and 561 lasers. A Plan Apochromat 60X (NA 1.4) oil immersion objective was used and the Fast Airyscan Detector was employed. Images were processed in Fiji (NIH).

### Endotoxemia *in vivo* model

Female wild-type and *Casp-11^-/-^* C57BL/6J mice between 8 and 10 weeks of age, were weighed and injected intraperitoneally 25 µg/g CL liposomes diluted in 10 mM HEPES (versus HEPES vehicle control). 10 minutes later, mice were injected with PBS or 10 µg/g of LPS from *Pseudomonas aeruginosa* (Sigma-Aldrich, L8643) diluted in PBS. 6 hours later, mice were weighed and humanely euthanised using CO2 and blood was collected by cardiac puncture. Blood was left to coagulate for 3 hours at RT, and serum was collected by centrifugation at 700 g for 10 minutes. Serum was assayed for circulating cytokines by ELISA.

### Quantification and statistical analysis

Statistical details of experiments, including the statistical tests used and the exact value of n, can be found in the figures and figure legends. For *in vitro* experiments, n represents the number of independent experiments (biological replicates). For *in vivo* experiments, n represents the number of mice per phenotype. In graphs, each symbol represents the mean value of technical triplicates from an independent experiment, while each bar represents the mean value across independent biological replicates, with the error bars representing the standard error of the mean (SEM) of three or more independent experiments, as indicated. Shapiro-Wilk tests were performed to assess whether data were normally distributed. All data were normally distributed, and the following parametric tests were performed: (1) unpaired t-test was used when comparing two independent groups (e.g., control vs. CL); (2) one-way analysis of variance (ANOVA) was used when comparing means among more than two groups (e.g., LPS vs LPS + CL vs LPS+RS-LPS) with Dunnett’s multiple-comparison test to compare all groups to a single control group (e.g., LPS + CL and LPS+RS-LPS compared to LPS), or Tukey’s multiple-comparison test to compare all pairs of groups (e.g., LPS vs LPS + CL and LPS vs LPS with inhibitors).

A significant p-value is indicated as *p<0.05; and p>0.05 as not significant (ns or not indicated). Nonlinear regression analysis was carried out using GraphPad Prism 10 software software (Graph-Pad Software, Inc.). Prism 10 was also used to generate graphs, calculate SEM and perform statistical analysis.

## Supplemental Information

The following supplementary figures are provided in the supplemental data file:

- Supplemental Figure S1: 18:2 CL does not induce cell death, and inhibits the noncanonical inflammasome regardless of the LPS origin
- Supplemental Figure S2: CL inhibits CASP11 activation independently of TLR4
- Supplemental Figure S3: CL inhibits the noncanonical inflammasome regardless of the transfection delivery sistem and accesses the cell interior to inhibit CASP11 activation independently of GBPs in BMDM

## Authors’ Contributions

Conceptualisation MP. Investigation MP, MM, SSB, JB, MK, JRC, EF, JCr, MDO, KMK, JCo. Methodology: MP, MM, SSB, JB, PP, KS. Validation MP, MM, SSB, JB, MK. Formal analysis MP, MM. Resources SMM, PB, PP and KS. Writing-original draft MP. Writing- Review and Editing MP, MM, SSB, JB, JCr, JCo, MK, SMM, PB, PP and KS. Visualisation: MP, MM. Funding acquisition: MP, MK, SMM, PB, PP and KS. Project administration: MP, SSB, PP and KS. Supervision: MP, SMM, PB, PP and KS.

## Acknowledgements

We gratefully acknowledge Dr James Curson and Prof Matthew Sweet (IMB, Brisbane, Australia) for providing BMDM from *Tlr4^-/-^* mice and Dr Madhavi Maddugoda (IMB, Brisbane, Australia) for editing this manuscript. We thank Maria del Carmen Baños and Ana Isabel Gomez (IMIB, Murcia, Spain) and Chinh Ngo (ANU, Canberra, Australia) for technical assistance with cell culture and plasmids.

