## Supplemental Data for "Cardiolipin Inhibits the Noncanonical Inflammasome by Preventing LPS Binding to Caspase 4/11 to Mitigate Endotoxemia in Vivo"

Pizzuto et al., 2024

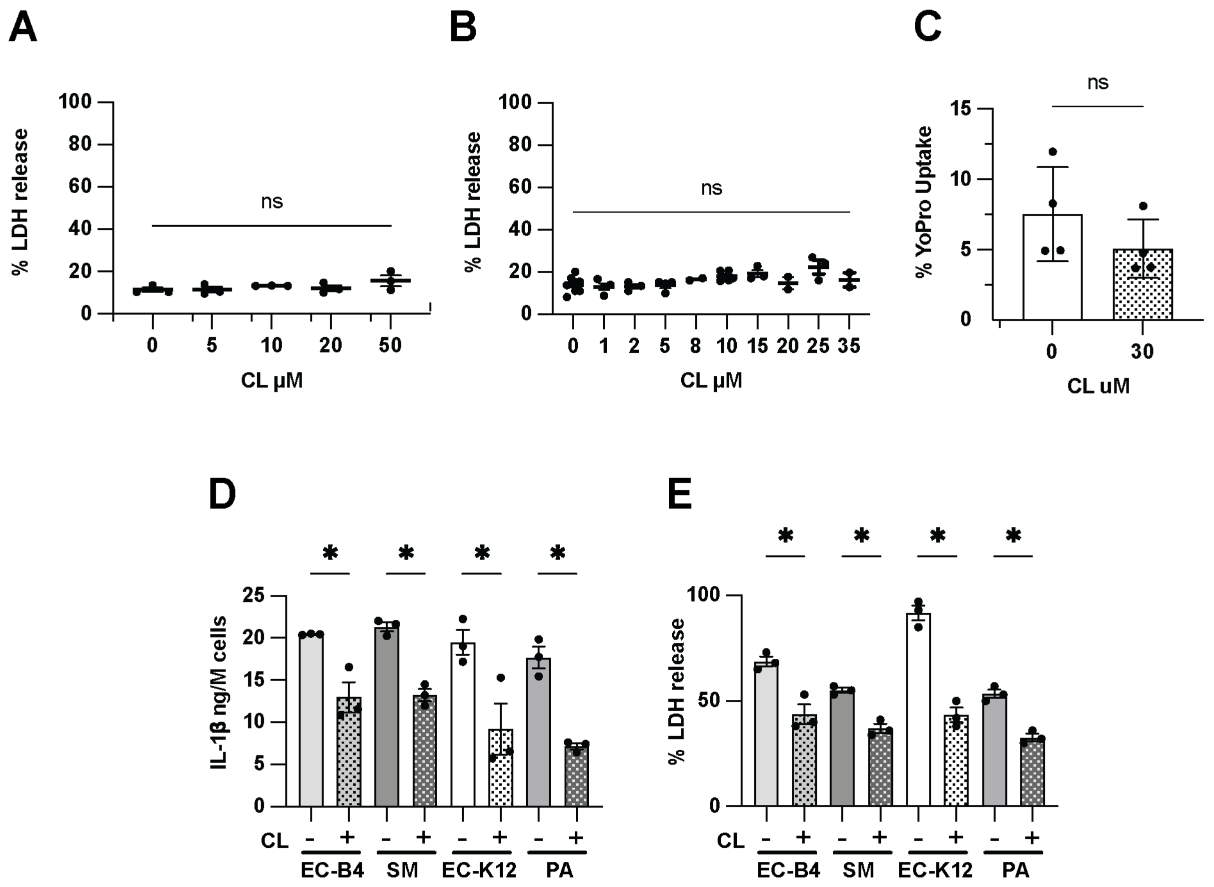


**Supplemental Figure S1: 18:2 CL does not induce cell death, and inhibits the noncanonical inflammasome regardless of the LPS origin**

**A – C** Human monocytes-derived macrophages (HMDM) from healthy donors (A) or bone marrow-derived macrophages (BMDM, B) from wild-type mice (WT) were incubated for 4 hours with 1 µg/mL $Pam_3CSK_4$. Cell culture medium was then replaced with OptiMEM (0) or increasing CL concentrations (1 to 50 µM) and cells were incubated for 18 hours. Lytic cell death was evaluated by LDH activity in supernatants (A-B) or YoPro uptake by lytic cells (C) and reported here as the percentage of cell lysis induced by 0.1% Triton.

**D – E** WT BMDM were incubated for 4 hours with 1 µg/mL $Pam_3CSK_4$. Cell culture medium was then replaced with OptiMEM plus 0.5% FuGENE HD complexed with LPS derived from EC-B4, *Escherichia coli* K12 strain (EC-K12), *Salmonella minnesota* R595 (SM), or *Pseudomonas aeruginosa* (PA) in the presence of HEPES (-) or 10 µM 18:2 CL. Cells were incubated for 18 hours. Cleaved IL-1β was quantified in cell supernatants by ELISA. LDH release was quantified by cytotoxicity assay.

**Data information:** Each symbol is the mean of three technical replicates from an independent biological replicate. Bars are the mean of three or more independent biological replicates (n = 3 to 4) ± SEM.

**Statistical analysis:** A and B one-way ANOVA compared to ctrl (0). C-I unpaired t-test. Significant difference for $p<0.05$ (*), not significant $p \geq 0.05$. Only comparisons of interest are shown.

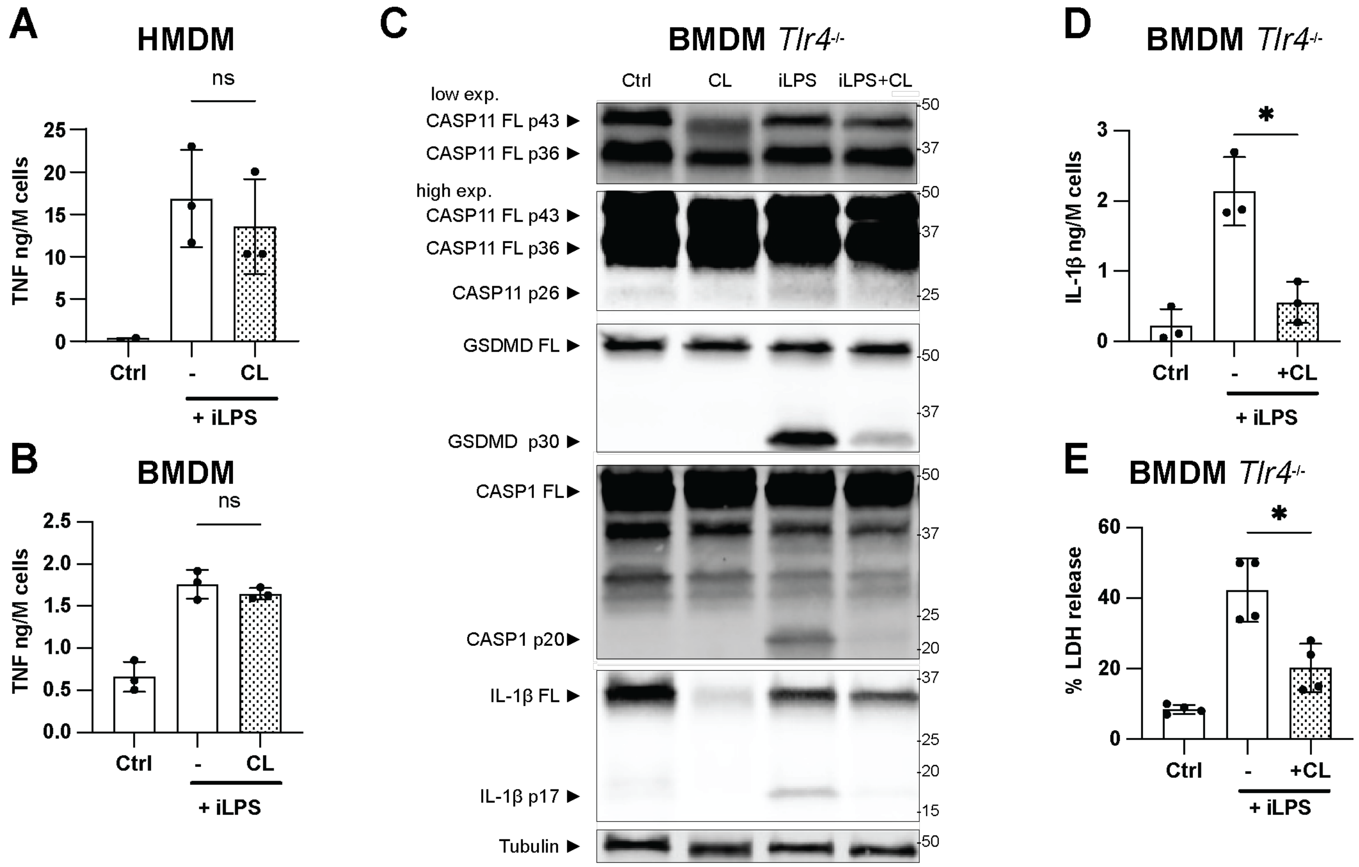


**Supplemental Figure S2: CL inhibits CASP11 activation independently of TLR4.**

HMDM from healthy donors (A), and BMDM from wild-type (B) or *Tlr4*$^{-/-}$ (C-E) mice were incubated for 4 hours with 1 µg/mL $Pam_3CSK_4$. Cell culture medium was then replaced with OptiMEM or the non-canonical inflammasome activator LPS complexed with 0.25 % v/v LTX (A) or 20 µg/mL CTB (B-E) (iLPS) in the presence of HEPES (-) or 10 µM CL. Released TNF and cleaved IL-1β were measured in supernatants by ELISA. CASP11, GSDMD, IL-1β and tubulin expression and cleavage were assessed by western blot. LDH release was quantified by cytotoxicity assay.

**Statistical analysis:** Unpaired t-test. Significant difference for p<0.05 (*), not significant p≥0.05. Only comparisons of interest are shown.

**Data information:** The blot is representative of three independent biological replicate experiments (n = 3). Each symbol is the mean of three technical replicates from an independent biological replicate. Bars are the mean of three or more independent biological replicates (n = 3 to 4) ± SEM.

**Statistical analysis:** Unpaired t-test. Significant difference for p<0.05 (*), not significant p≥0.05 (ns).

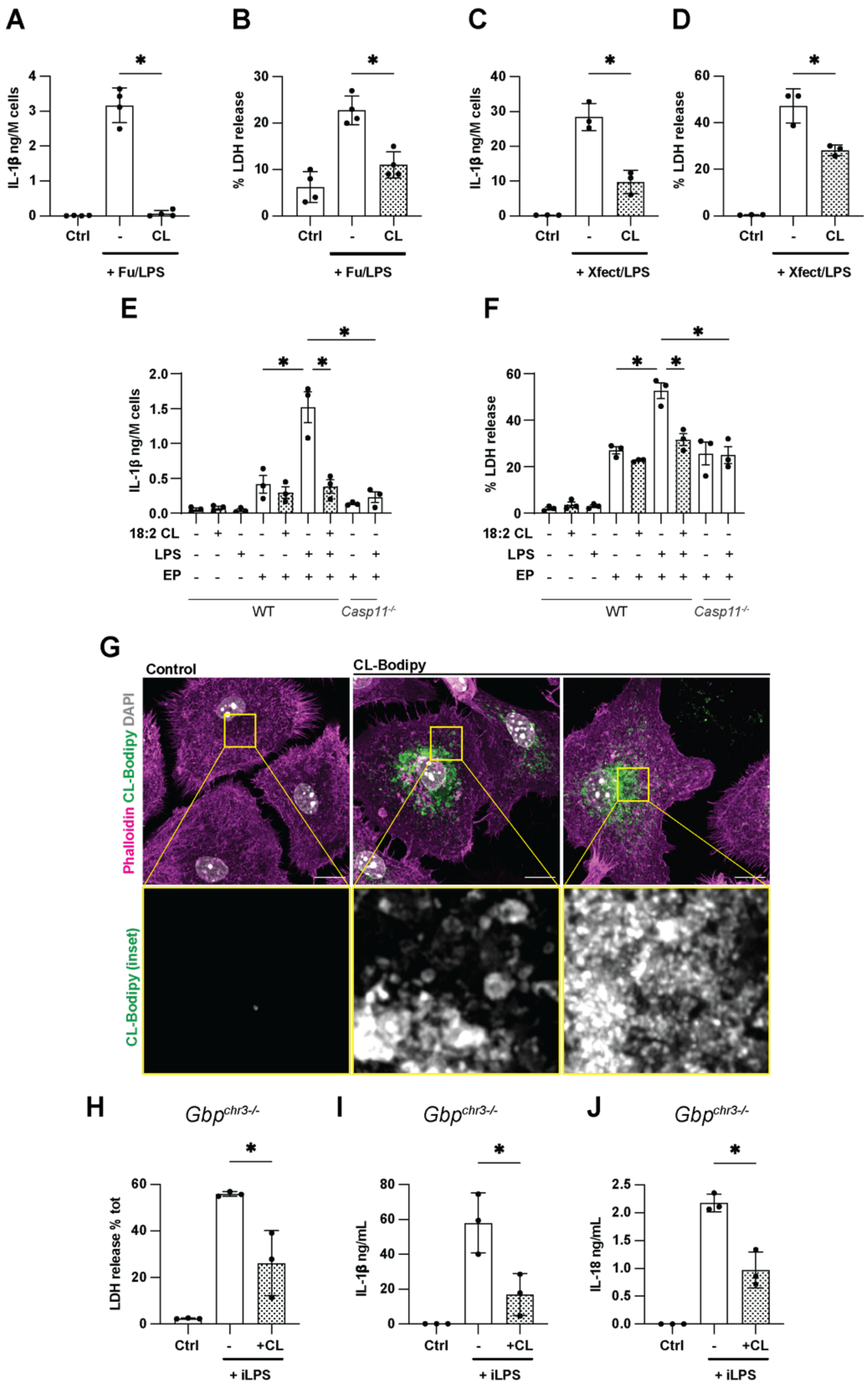

A
B
C
D
IL-1β ng/M cells
% LDH release
Ctrl
-
CL
+ Fu/LPS
+ Xfect/LPS
E
F
18:2 CL
LPS
EP
WT
Casp11-/-
G
Control
CL-Bodipy
Phalloidin CL-Bodipy DAPI
CL-Bodipy (inset)
H
I
J
Gbpchr3-/-
LDH release % tot
IL-1β ng/mL
IL-18 ng/mL
+CL
+ iLPS

**Supplemental Figure S3: CL inhibits the noncanonical inflammasome regardless of the delivery system used for LPS, and accesses the cell interior to inhibit CASP11 activation independently of GBPs in BMDM**

**A-D** BMDM from WT mice were incubated for 4 hours with 1 µg/mL $Pam_3CSK_4$. Cell culture medium was then replaced with OptiMEM (Ctrl and -), or 2 µg/mL of the noncanonical inflammasome activator LPS from *Escherichia coli* B4 strain (EC-B4) complexed with 0.5% FuGENE HD (A-B) or 1.2% Xfect (C-D) (iLPS) in the presence of HEPES (-) or 10 µM CL. Cells were incubated for 18 (A-B) or 4 hours (C-D). Cleaved IL-1β was quantified in cell supernatants by ELISA. LDH release was quantified by cytotoxicity assay.

**E-F** BMDM from wild-type or *Casp11*$^{-/-}$ mice were incubated for 4 hours with 1 µg/mL $Pam_3CSK_4$. Cell culture medium was then replaced with HEPES-supplemented OptiMEM in the presence of HEPES (-,10 µM CL, or 2 µg/mL of LPS. Cells were left untouched or electroporated (+EP), then fresh OptiMEM was added, and cells were incubated for 4 hours. Cleaved IL-1β was quantified in cell supernatants by ELISA. LDH release was quantified by cytotoxicity assay.

**G** Fixed-Airyscan confocal imaging of wild-type BMDM primed for 4 hours with 1 µg/mL $Pam_3CSK_4$, and incubated with HEPES (Ctrl) or 10 µM of TopFluor CL (BODIPY-CL) for a further 18 hours. Macrophages were stained with the actin probe phalloidin (magenta), BODIPY-CL (green) and DAPI (grey). Images are maximum intensity projections of Z-stack acquisitions. Scale bar= 10 µm.

**H-J** BMDM from *Gbp*$^{ch3-/-}$ mice were incubated for 4 hours with 1 µg/mL $Pam_3CSK_4$. Cell culture medium was then replaced with OptiMEM (ctrl) or LPS complexed with 1.2% Xfect in the presence of HEPES (-) or 10 µM 18:2 CL. Cells were incubated for 4 hours. LDH release was quantified by cytotoxicity assay (H). Cleaved IL-1β (I) and total IL-18 (J) were quantified in cell supernatants by ELISA.

**Statistical analysis:** A-B One-way ANOVA Dunnett's multiple comparisons test. D-F Unpaired t-test. Significant difference for $p<0.05$ (*), not significant $p\geq0.05$. Only comparisons of interest are shown.

**Data information:** Images are representative of three independent biological replicates (n = 3). Each symbol is the mean of technical triplicates from an independent biological replicate. Bars are the mean of three independent biological replicates (n = 3) ± SEM.

**Statistical analysis:** Unpaired t-test. Significant difference for $p<0.05$ (*), not significant $p\geq0.05$ (ns).
